# A novel tumor-targeted interferon-alpha/-beta receptor 1 antagonist increases replication of oncolytic vesicular stomatitis virus in a mouse mesothelioma model

**DOI:** 10.64898/2026.06.02.729496

**Authors:** Thomas Teja Ogor, Yann Bordat, Manon Souchard, Joëlle Nader, Geneviève Garcin, Camille Chatelain, Virginie Dehame, Sophie Deshayes, Lucas Treps, Maria Naranjo-Gomez, Nicolas Boisgerault, Jan Tavernier, Mireia Pelegrin, Jean-François Fonteneau

**Affiliations:** Université de Montpellier, LPHI, 34000 Montpellier, France; Nantes Université, INSERM UMR 1307, CNRS UMR 6075, Université d’Angers, CRCI2NA, F-44000 Nantes, France; Labex IGO, Immunology Graft Oncology, 44007 Nantes, France; IRMB, Univ Montpellier, INSERM, CNRS, Montpellier, France; Labex Mabimprove, Tours-Montpellier, France; Cytokine Receptor Laboratory, VIB Medical Biotechnology Center, Ghent University, 9000 Ghent, Belgium

**Keywords:** Type I interferons, interferon-alpha/-beta receptor, interferon-stimulated genes, bispecific nanobodies conjugates, cell-specific IFNAR antagonist, oncolytic virus, vesicular stomatitis virus, tumor virotherapy

## Abstract

Type I Interferons (IFN-I) are cytokines with pleiotropic activities involved in antiviral and antitumor immune responses. They can reduce oncolytic virus replication in tumor cells by inducing expression of interferon stimulated genes (ISG) with antiviral functions. To specifically neutralize the IFN-α/-β receptor (IFNAR) on specific cell types, we created novel IFNAR1-targeted antagonists constituted of a high-affinity nanobody targeting a specific cell surface marker conjugated to a low-affinity blocking nanobody targeting IFNAR1. We first show *in vitro* and *in vivo* that such an antagonist targeting the mouse CD20 molecule (mCD20) inhibits IFNAR signaling only in B cells among splenocytes. We then showed *in vitro* that a human CD20 (hCD20)-targeted antagonist blocks IFNAR signaling and induces vesicular stomatitis virus (VSV) oncolytic activity against IFN-α11-treated AK7 mesothelioma or B16 melanoma cells only if these cells express hCD20. *In vivo*, we show that the antagonist binds to hCD20 and enhances VSV replication by inhibiting ISG expression specifically in hCD20+ AK7 mesothelioma tumors. Altogether our results demonstrate the efficient and cell-type specific inhibition of IFNAR signaling through the use of these novel IFNAR1 antagonists, both *in vitro* and *in vivo*. These antagonists could have many therapeutic applications given the importance of IFN-I in numerous diseases.

**Graphical abstract:** 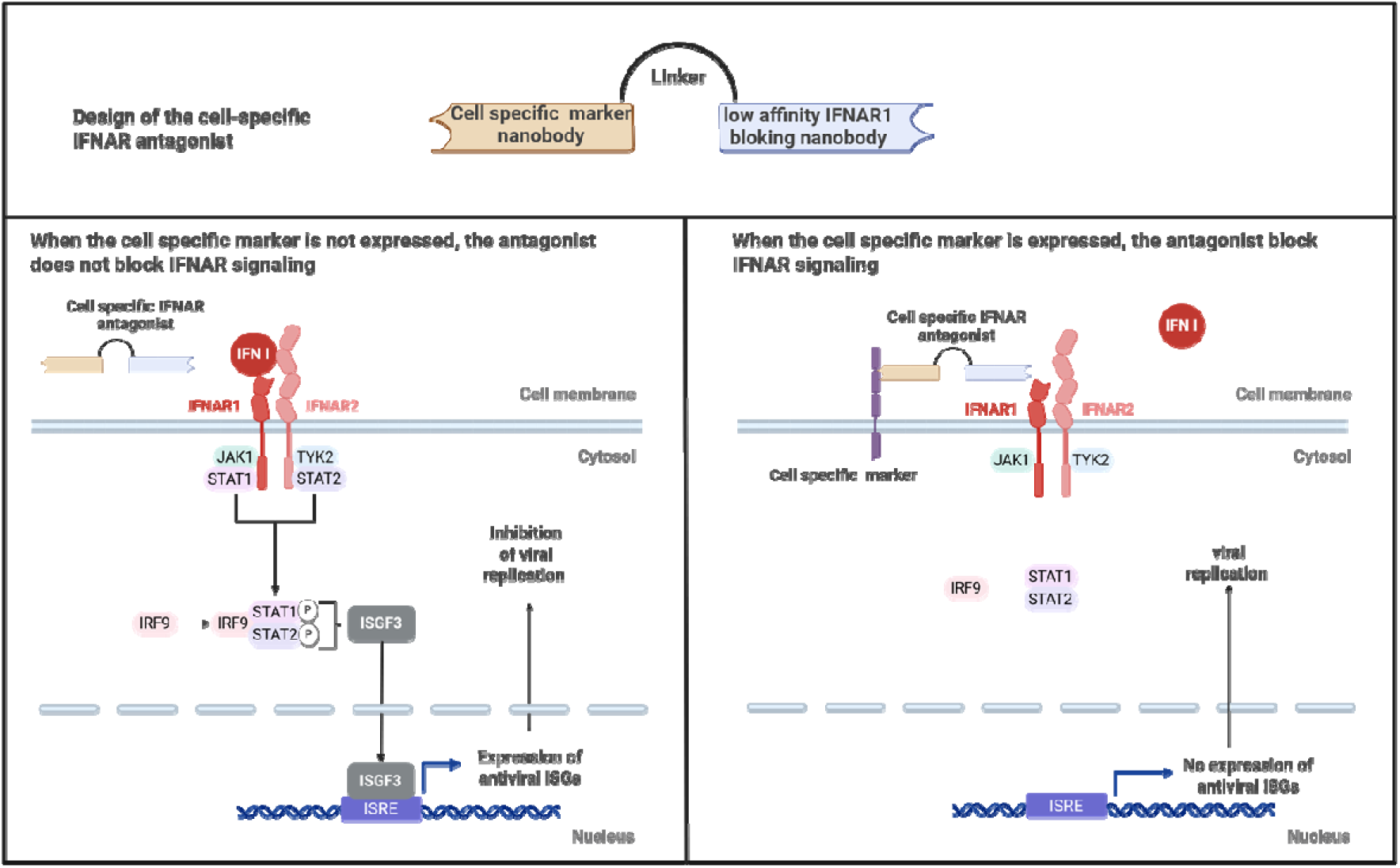

**eTOC synopsis:** In this study, we created cell-specific IFNAR antagonists that allow to inhibit selectively IFNAR signaling in particular types of cell. We show that this antagonist can be used to target IFNAR at the surface of tumor cells that lead to the inhibition of IFNAR signaling and ISG expression in these cells rendering them more permissive to VSV replication. Beside antitumor virotherapy, these novel antagonist could be useful to study role of IFN-I in normal or pathological context.

## Introduction

The type I interferon (IFN-I) response allows infected cells to report *via* IFN-α and IFN-β secretion the presence of viruses to neighboring cells and to the immune system ^1^. Infected cells are able to detect viral genomes in their cytoplasm and produce IFN-I in response *via* the stimulator of interferon genes protein (STING) pathway for DNA viruses and the mitochondrial antiviral signaling protein (MAVS) pathway for RNA viruses. The interferon-alpha/-beta receptor (IFNAR), composed of the subunits IFNAR1 and IFNAR2, allows cells exposed to IFN-I to express hundreds of IFN-stimulated genes (ISG) that induce a state of antiviral resistance. Many ISG act directly by blocking the different stages of the viral cycle, from the entry of the virus, through the inhibition of its replication, to the release of its progeny by the infected cell. The IFN-I response also plays a crucial role in the induction and the regulation of the adaptive antiviral immune response, notably by favoring antigen cross-priming by dendritic cells ^2,3^.

Oncolytic immunotherapy is a therapeutic approach based on oncolytic viruses (OVs) ^4^. OVs are non-pathogenic viruses that preferentially replicate in and kill tumor cells without harming healthy cells. Furthermore, replication of OVs in tumor cells leads to immunogenic death that can induce immune cell infiltration of tumors to reinforce the antitumor response, notably through the induction of the IFN-I response^5^.

Some OVs such as attenuated strains of measles virus (MV) or vesicular stomatitis virus (VSV) are highly sensitive to the antiviral properties of IFNAR signaling ^6–9^. In tumor cells with defects of IFNAR signaling, IFN-I are not able to block OV replication ^6^, whereas in tumor cells without defect of IFNAR signaling, IFN-I may limit viral replication and thus oncolytic activity ^7,8,10^. Furthermore, IFN-I response is often activated in tumors, as reported by an analysis of 31 cancers *via* The Cancer Genome Atlas (TCGA) database by Liu H et al who analyzed the intensity of an IFN-I signature based on the expression of 38 ISG ^11^. This constitutive IFN-I response found in some tumors has been shown to limit oncolytic efficacy ^9^.

Thus, IFN-I response is a double-edged sword for oncolytic immunotherapy. On one hand, the induction of IFN-I response by OVs inflames the tumor environment leading to an enhanced adaptive antitumor immune response. Furthermore, IFN-I protect healthy tissue from viral replication. On the other hand, a pre-existing or an OV-induced IFN-I response may quickly shut down the replication and spreading of OVs inside tumors. Therefore, it would be interesting to design a strategy to transiently block IFNAR signaling and thus ISG expression only in tumor cells during OV treatment. This strategy would enhance viral replication in tumor cells, while preserving the positive effects of IFN-I on the protection of healthy tissues and the induction of the antitumor immune response.

Recently, we reported a strategy to induce IFNAR signaling only in a particular cell type ^12^. This strategy consists in a mutated IFN-α2 with a lower affinity to IFNAR compared to natural IFN-α2. This mutated IFN-α2 is then conjugated to a single variable domain of heavy chain antibodies (VHHs), also known as nanobody which recognizes specifically a marker expressed by the targeted cell type. Using a mutated IFN-α2 conjugated to a nanobody that recognizes Clec9a, we showed that this targeted agonist induced *in vitro* and *in vivo* IFNAR signaling only in CD8+ CD11c+ dendritic cells that also expressed Clec9a ^13^.

In the present study, we developed a similar strategy to target and block IFNAR signaling in particular cell types using a low-affinity anti-IFNAR1 antagonist nanobody. Thus, we designed a cell-targeted IFNAR1 antagonist by coupling the anti-IFNAR1 nanobody to a high-affinity nanobody recognizing a specific cell surface marker to inhibit IFNAR signaling only when the second nanobody is bound to its target. Using a murine CD20 (mCD20)-targeted IFNAR1 antagonist, we first show that our strategy is able to block IFNAR signaling induced by an intravenous injection of IFN-I, only in CD19+ splenic B lymphocytes. We then show that an hCD20-targeted IFNAR antagonist similarly reduces IFNAR signaling and ISG expression in tumor cells, selectively enhances intratumor VSV replication and modestly increases its oncolytic activity in the orthotopic hCD20+ AK7 peritoneal mesothelioma mouse model. Overall, our results demonstrate the selective blocking of IFNAR signaling with a novel nanobody-based IFNAR1 antagonist.

## Results

### Designing cell-targeted IFNAR1 antagonists

We first generated recombinant IFNAR1 antagonist nanobodies capable of neutralizing the IFNAR signaling with low affinity. Llamas were immunized against the murine IFNAR1 protein to obtain a library of 61 VHH candidates with 20 different CDR3 region groups, of which 10 were deemed neutralizing (Figure 1A). Four IFNAR1 neutralizing VHH candidates were then fused, through a 20X Gly-Gly-Ser (GGS) linker, with a second VHH that targets with high affinity the mouse CD20 (mCD20) or the human CD20 (hCD20) surface markers ^14^. The capacity of the four mCD20-targeted IFNAR1 antagonists to block IFNAR signaling, and thus STAT1 phosphorylation, specifically in CD19+CD20+ B cells was measured *in vitro* on splenocytes exposed to IFN-I (Figure 1B-D). CD19 was used to discriminate B cells since CD20 is masked by the antagonist. This allowed us to obtain two targeted antagonists with high targeting potential: candidate-3 and candidate-4, which displayed IC₅₀ values that were 1,447- and 3,444-fold lower on mCD20⁺ cells than on mCD20⁻ cells, respectively. For the rest of the study, we used the mCD20- and hCD20-targeted IFNAR1 antagonists obtained with candidate-3, since this nanobody has a lower IC_50_ (0.06nM) than candidate-4 (0.34nM) on targeted cells. For *in vivo* studies, the 20X GGS linker of this construct was replaced with a 300 amino acids PASylated linker to increase the antagonist half-life when injected into mice ^15^.

**Figure 1:**
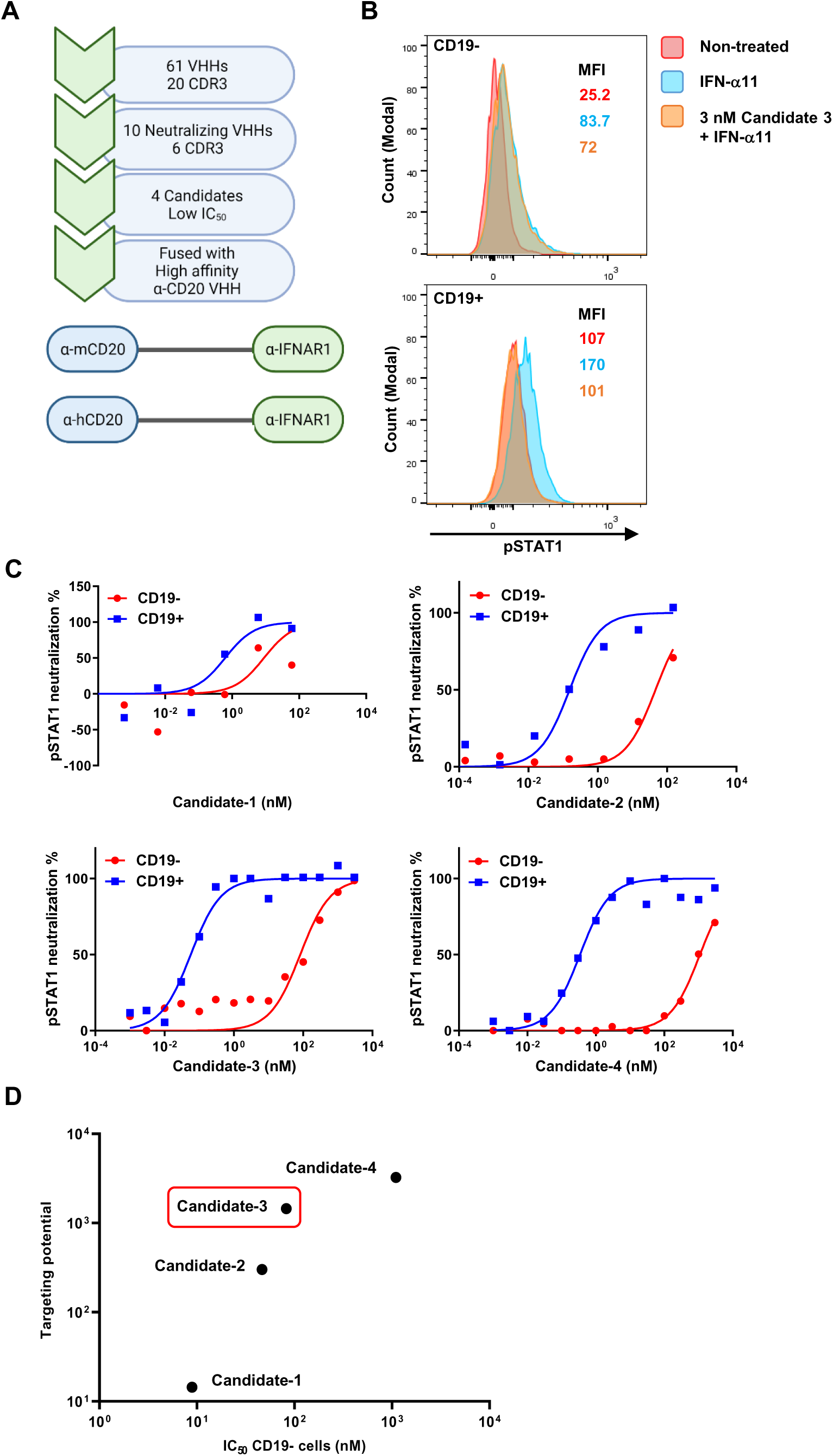
mCD20-targeted IFNAR1 antagonist blocks IFNAR signaling in B cells *in vitro*. A) In order to specifically target the IFNAR signaling on different types of cells, we obtained a library of anti-IFNAR1 VHHs through the immunization of lamas and subsequent phage display screening of the obtained antibodies library. From this library of anti-IFNAR1 VHHs, four low affinity neutralizing candidates were chosen and fused with a high affinity VHH targeting either human or murine CD20. B) Mouse splenocytes isolated from Balb/c mice were exposed *in vitro* to IFN-α11 alone or to IFN-α11 in combination with the candidate-3 mCD20-targeted IFNAR1 antagonist during 30 minutes. The induction of pSTAT1 was measured by flow cytometry. C) Mouse splenocytes were exposed *in vitro* to IFN-α11 alone or to IFNα-11 in combination with increasing concentrations of the four mCD20-targeted IFNAR1 antagonist candidates during 30 minutes. The induction of pSTAT1 was measured by flow cytometry. The % of pSTAT1 neutralization was calculated as following: ((pStat1 MFI of cells treated with IFN-α11 - pStat1 MFI of cells treated with IFN-α11 and the antagonist) / (pStat1 MFI of cells treated with IFN-α11 - pStat1 MFI of untreated cells)) x 100. MFI represents the mean of fluorescence. D) The targeting potential of the four candidates fused with the nanobody against mCD20 was calculated as the ratio of the IC_50_ between non-targeted (CD19-CD20-) and targeted (CD19+CD20+) splenocytes.

### The mCD20-targeted IFNAR1 antagonist neutralizes IFNAR signaling only in B cells in vivo

After having demonstrated the functional effect of this B cell-targeted IFNAR1 antagonist *in vitro*, we sought to test its capabilities *in vivo*. Mice were co-injected intravenously (i.v.) with a natural mix of IFN-I five minutes after the mCD20-targeted IFNAR1 antagonist to inhibit IFNAR signaling only in murine CD19+CD20+ B cells (Figure 2A). The mix of IFN-I was recovered from infected cells and contains several IFN-α and the IFN-β. It allows to evaluate the antagonist effect against a combination of different IFN-I. We then assessed the expression of ISG54 and PD-L1 on collected splenocytes by flow cytometry (Figure 2B and 2C). Both proteins are coded by ISG and therefore serve as reliable indicators of IFNAR signaling. Results showed that CD19+ B cells representing 35% of total splenocytes did not express PD-L1 nor ISG54 when receiving both IFN-I and the mCD20-targeted IFNAR1 antagonist, whereas all CD19- splenocytes expressed both proteins at levels comparable to those observed in IFN-I-treated control mice. The mCD20-targeted IFNAR1 antagonist can thus efficiently and specifically inhibit IFNAR signaling in B cells *in vivo*.

**Figure 2:**
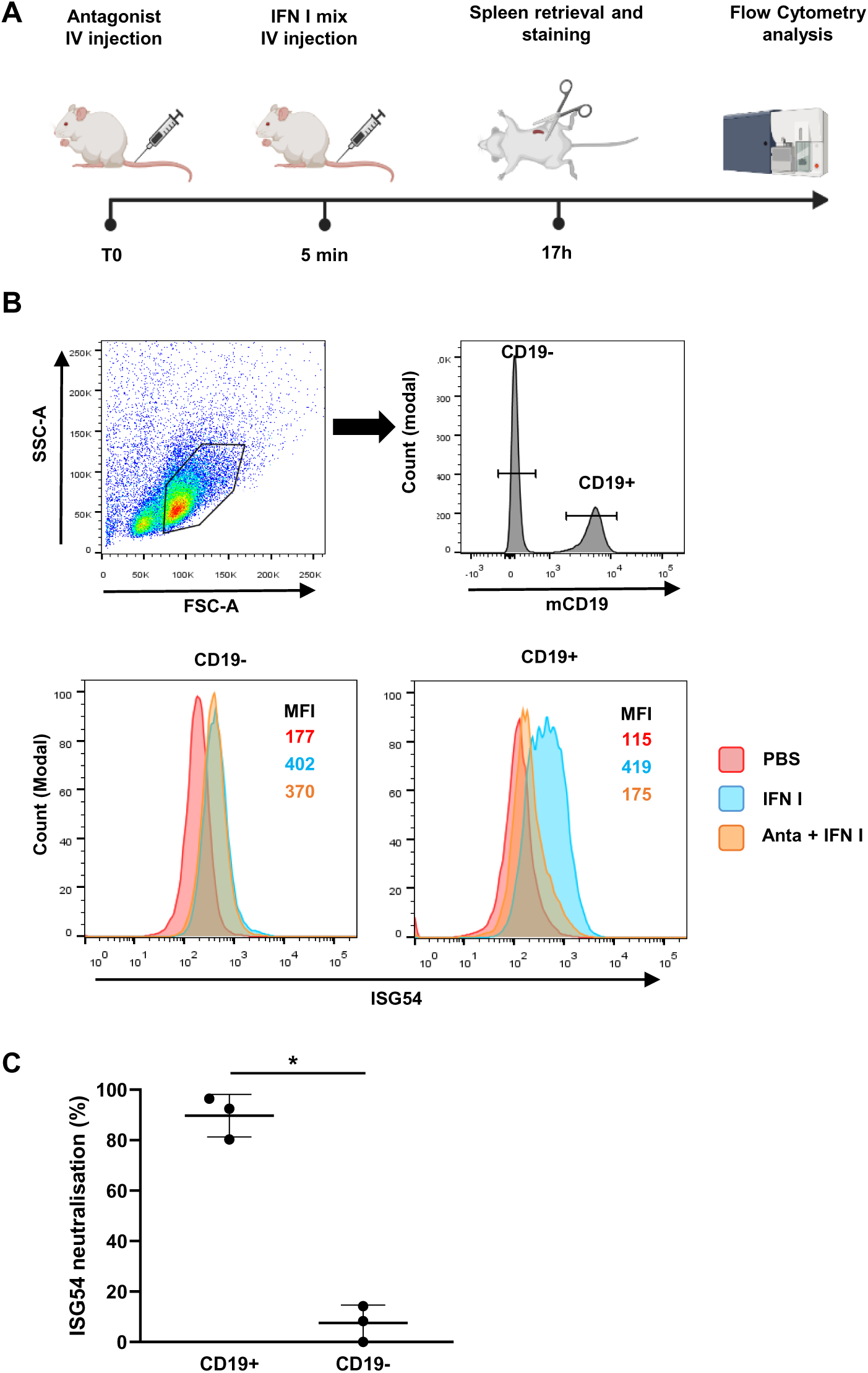
the mCD20-targeted IFNAR1 antagonist blocks IFNAR signaling in B cells *in vivo*. A) Time line of the experiment. B) C57/BL6 mice received PBS, 30000 U of a natural mix of IFN-I or a natural mix of IFN-I and 10 μg of the mCD20-targeted IFNAR1 antagonist. After 17h, splenocytes were isolated and their CD19 and ISG54 expression was analyzed by flow cytometry. C) Neutralization % of ISG54 expression was calculated by dividing the difference of MFI between the IFN-I control and the antagonist-treated samples, to the difference of MFIs between the IFN-I and PBS controls, and multiplied by 100. Results were represented as means ± SEM of four different experiments.

We then confirmed that our approach can be used to target other cell subsets such as Clec9A+ dendritic cells (DC) (Supplemental Figure 1). Clec9A+ DC can be discriminated among the splenocytes by the co-expression of CD8 and CD11c. We showed *in vivo* that after IFN-I injection a Clec9A-targeted IFNAR1 antagonist was capable of blocking IFNAR signaling only in CD8+CD11c+ DC among splenocytes.

### The hCD20-targeted IFNAR1 antagonist is able to neutralize IFNAR signaling in hCD20+ mouse tumor cell lines and induce their permissiveness to VSV replication

Having validated *in vivo* the cell-specific targeting potential of the IFNAR1 antagonists on different mouse splenocyte subsets, we sought to test this IFN-I neutralization approach on murine tumor cell lines. Our strategy relies on the successful recognition of a tumor antigen in order to specifically neutralize IFN-I activity on tumor cells while sparing IFNAR from healthy tissues. We chose to target a surface marker of human origin excluding the possibility for the targeting moiety to bind to mouse cells *in vivo*. To this end, we engineered a recombinant IFNAR1 antagonist that binds to the hCD20 protein and tested its binding capacity on the mesothelioma AK7 and melanoma B16 cell lines transfected with a plasmid encoding hCD20. Both cell lines strongly expressed this marker on their surface (Supplemental Figure 2).

The next step was to assess whether the hCD20-targeted IFNAR antagonist could neutralize the activity of IFN-I in both hCD20+ cancer cell lines. As demonstrated in Figure 3A and 3B, the hCD20-targeted IFNAR1 antagonist was indeed able to neutralize, in a dose-dependent manner, the expression of ISG54 and PD-L1 specifically on the hCD20+ AK7 and hCD20+ B16 cancer cell lines with an IC_50_ inferior to 1 nM. In contrast, this neutralization was not observed in the hCD20-AK7 and mCD20+ B16 control cell lines, demonstrating the hCD20 specific inhibition of IFNAR signaling by the antagonist.

**Figure 3:**
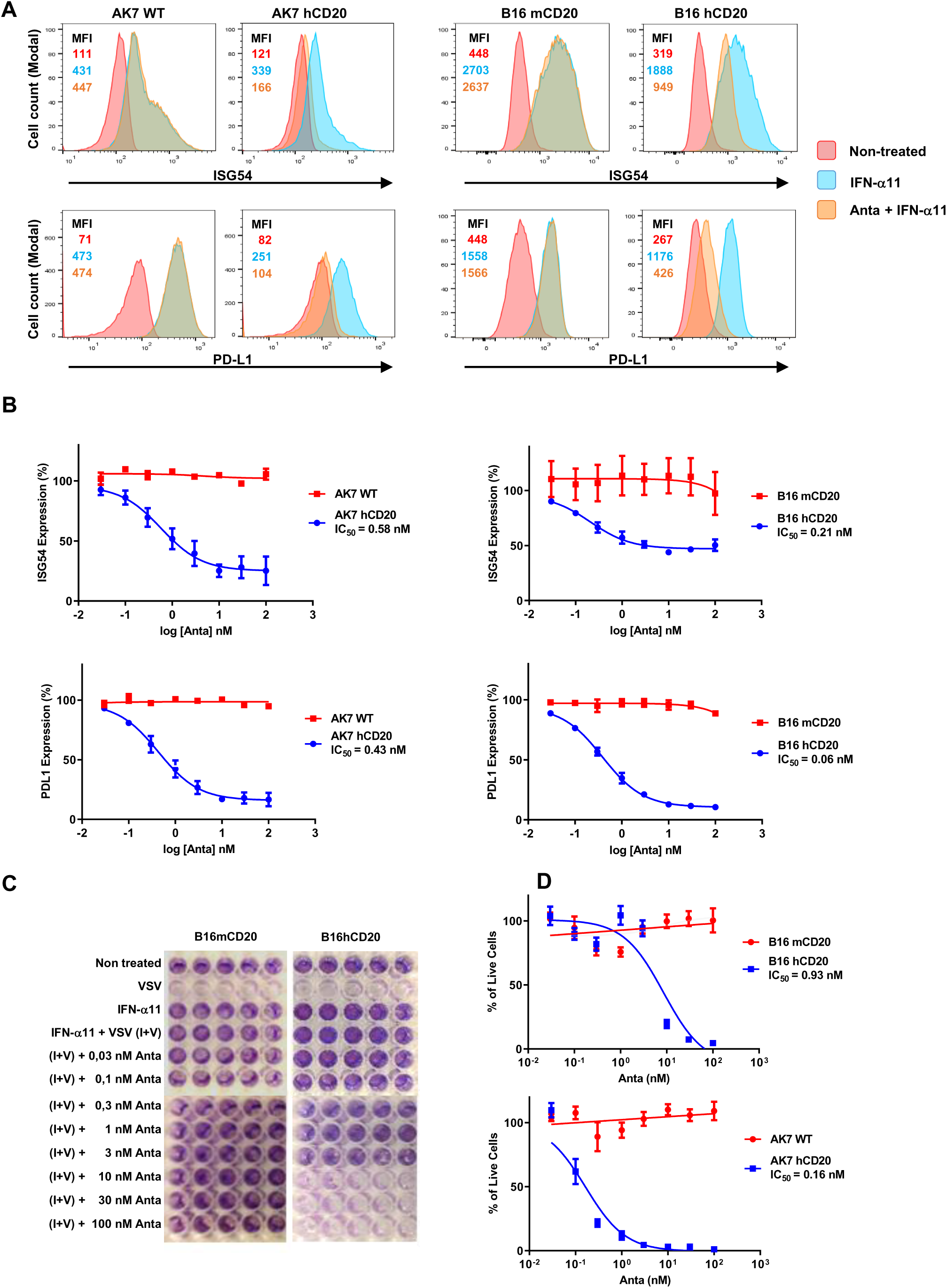
the hCD20-targeted IFNAR1 antagonist blocks IFNAR signaling specifically on hCD20+ B16 melanoma and hCD20+ AK7 mesothelioma cells. A) hCD20- or hCD20+ AK7 cancer cell lines were treated either with 10 nM of hCD20-targeted IFNAR antagonist and IFN-α11 (3000 U/ml), with IFN-α11 (3000 U/ml) alone or left untreated. Antagonist was added 30 minutes before IFN-α11. 24h later, ISG54 expression was measured by flow cytometry. B) hCD20- or hCD20+ B16 or AK7 cell lines were treated with IFN-α11 in presence of increasing doses of hCD20-targeted IFNAR antagonist. 24h later, ISG54 and PDL1 expression was measured by flow cytometry. Neutralization % was calculated by dividing the difference of MFIs between the IFN-I-treated and the IFN-I/antagonist-treated samples, to the difference of MFIs between the IFN-I-treated samples and PBS-treated controls, and multiplying by 100. Results were represented as means ± SEM of three different independent experiments. C) After treatment with different doses of hCD20-targeted IFNAR antagonist, B16mCD20 or B16hCD20 tumor cells were exposed to 100 U/ml IFN-α11 alone or to 100 U/ml IFN-α11 and VSV (I+V) (MOI=0,12). Cells were stained and fixed with a violet crystal solution to identify their viability. D) After 48h hours exposition to hCD20-targeted IFNAR1 antagonist, 100 U/ml of IFN-α11 and/or 5.7×10^4^ PFU/ml of VSV, live cells identified by violet crystal staining were quantified. Live cells % was calculated by dividing the difference of absorbance at 570 nm between the IFN-I-treated and the IFN-I/antagonist-treated samples, by the difference of absorbance at 570 nm between the IFN-I-treated and non-treated samples, and multiplying by 100. Results were represented as means ± SEM of three different experiments.

One of the main concerns for effective cancer virotherapy is the possible sensitivity of tumor cells to the antiviral effects of IFN-I that reduces viral replication and oncolytic activity. Thus, we wanted to determine if the hCD20-targeted IFNAR antagonist was able to increase the viral oncolytic activity only on hCD20+ tumor cells. As an OV, we used VSV, a rhabdovirus that harbors an extremely low toxicity in IFN-I-competent experimental models ^16,17^.

IFN-α11-treated mCD20+ B16 cells were completely protected from the oncolytic activity of VSV in absence or presence of the hCD20-targeted IFNAR1 antagonist (Figure 3C). The antagonist specifically neutralized the antiviral properties of IFN-α11 only in hCD20+ B16 tumor cells at relatively low concentrations. Indeed, hCD20+ B16 or AK7 cell lines that were treated with as little as 10 nM or 3 nM of hCD20-targeted IFNAR1 antagonist, respectively, were entirely killed by VSV 48h post-infection (Figure 3D). The neutralizing action of the antagonist was not observed in B16 or AK7 cells that do not express hCD20. This shows that the hCD20-targeted IFNAR1 antagonist is able to block IFNAR signaling and to specifically sensitize hCD20+ tumor cells to VSV oncolytic activity.

### VSV injection in mice induces IFN-I secretion and IFNAR signaling in tumor and spleen

To assess the induction of IFN-I secretion and ISG expression *in vivo* after VSV injection, we used the syngeneic intraperitoneal AK7 mesothelioma model in C57BL/6 female mice ^18^. This tumor model is characterized by the early colonization of the omentum and the formation of cancer cell aggregates suspended in the peritoneal cavity. Late stages of the disease see the invasion of other connective tissues such as the mesentery, and the formation of peritoneal ascites, at which point mice are euthanized.

Based on the observation of ISG expression in the tumors of untreated animals 7 days post-implantation (data not shown), we chose to work with early tumors at 3 days post-implantation, in order to observe ISG expression induced by VSV. As soon as 5h after injection of 1.5×10^7^ PFU of VSV through the caudal vein, IFN-α could be detected in the serum (Figure 4A). This IFN-I production rapidly induced ISG expression in the spleen and later in the tumor (Figure 4B). These results suggest that the hCD20-targeted IFNAR antagonist should be administered at the time of VSV injection in order to block IFNAR signaling in tumor cells and consequently enhance VSV oncolytic activity.

**Figure 4:**
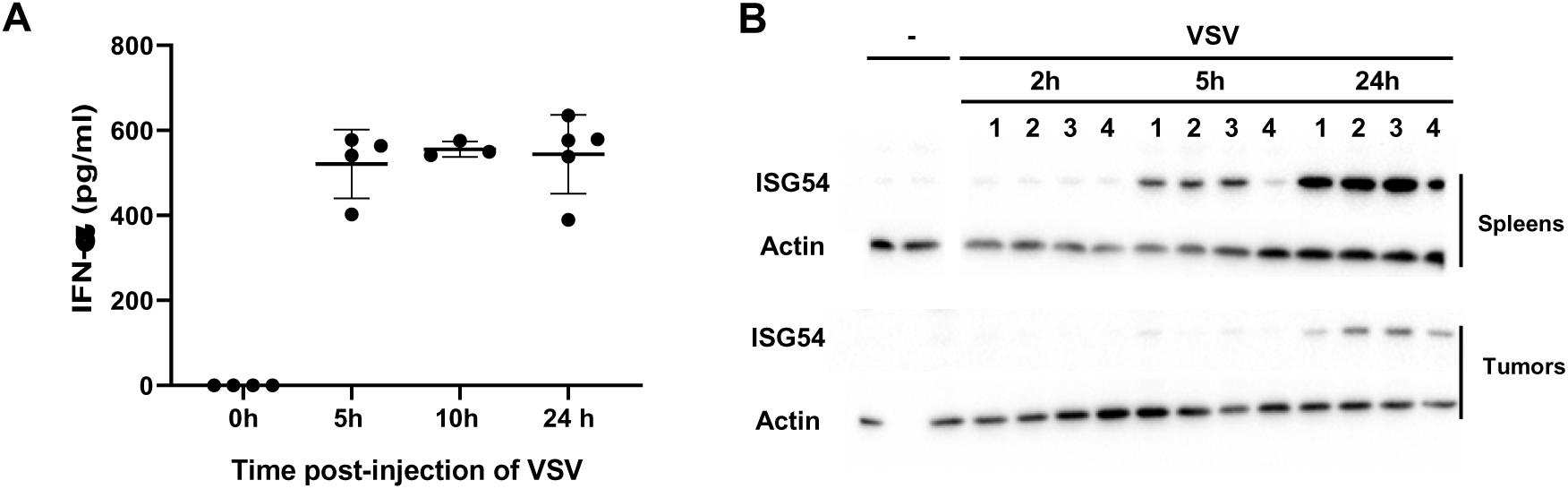
infection by VSV triggers the IFN-I response and ISG expression in AK7 tumor bearing mice. A) AK7 bearing mice were injected intravenously with 1.5×10^7^ PFU of VSV. At different time points after injection, blood was collected and IFN-α measured by ELISA. Results are the means ± SEM of three independent experiments. B) AK7 bearing mice were injected with 1.5×10^7^ PFU of VSV. At different time points after injection, tumors and spleens were collected from 4 mice to measure ISG54 expression by western blot in each of them.

### The hCD20-targeted IFNAR1 antagonist enhances VSV replication in hCD20+ AK7 tumor in vivo

We next determined if the hCD20-targeted IFNAR1 antagonist blocks IFNAR signaling and enhances VSV replication in hCD20+ tumors *in vivo.* Mice bearing hCD20+ AK7 tumors received 1.5×10^7^ PFU of VSV alone or combined with two injections of 10 µg of the antagonist at the time of VSV administration and 12h later (Figure 5A). 36h after VSV injection, tumors were dissociated. We observed that tumors treated with the antagonist lost detectable expression of hCD20, suggesting that the antagonist was already bound to hCD20 (Figure 5B). We then assessed the functional activity of the tumor-targeted IFNAR1 antagonist by analyzing the expression of cellular ISG54 and two viral genes, *VSV-G* and *VSV-N*, in the tumors and spleens of treated animals. In the absence of antagonist, ISG54 was detected in CD45- tumor cells and CD45+ non-malignant cells from the tumor microenvironment (Figure 5C). ISG54 expression was blocked by the hCD20-targeted IFNAR1 antagonist in CD45- cells, whereas no blocking was observed in CD45+ cells. At 36 hours post-treatment, expression levels of VSV-G and VSV-N were increased by 76.6-fold and 50.6-fold, respectively, respectively, in hCD20+ AK7 tumors from mice that received the VSV/antagonist combination compared to mice that received the VSV alone (Figure 5D). We also observed a smaller increase of VSV-G (15.3-fold) and VSV-N (9.9-fold) expression in the spleens of animals that received the combination. Altogether these results indicate that the use of an hCD20-targeted IFNAR1 antagonist enhances VSV replication in hCD20+ AK7 tumors, but also suggest that the increased replication of VSV in tumors may spill over in the mouse circulation, reaching the spleen.

**Figure 5:**
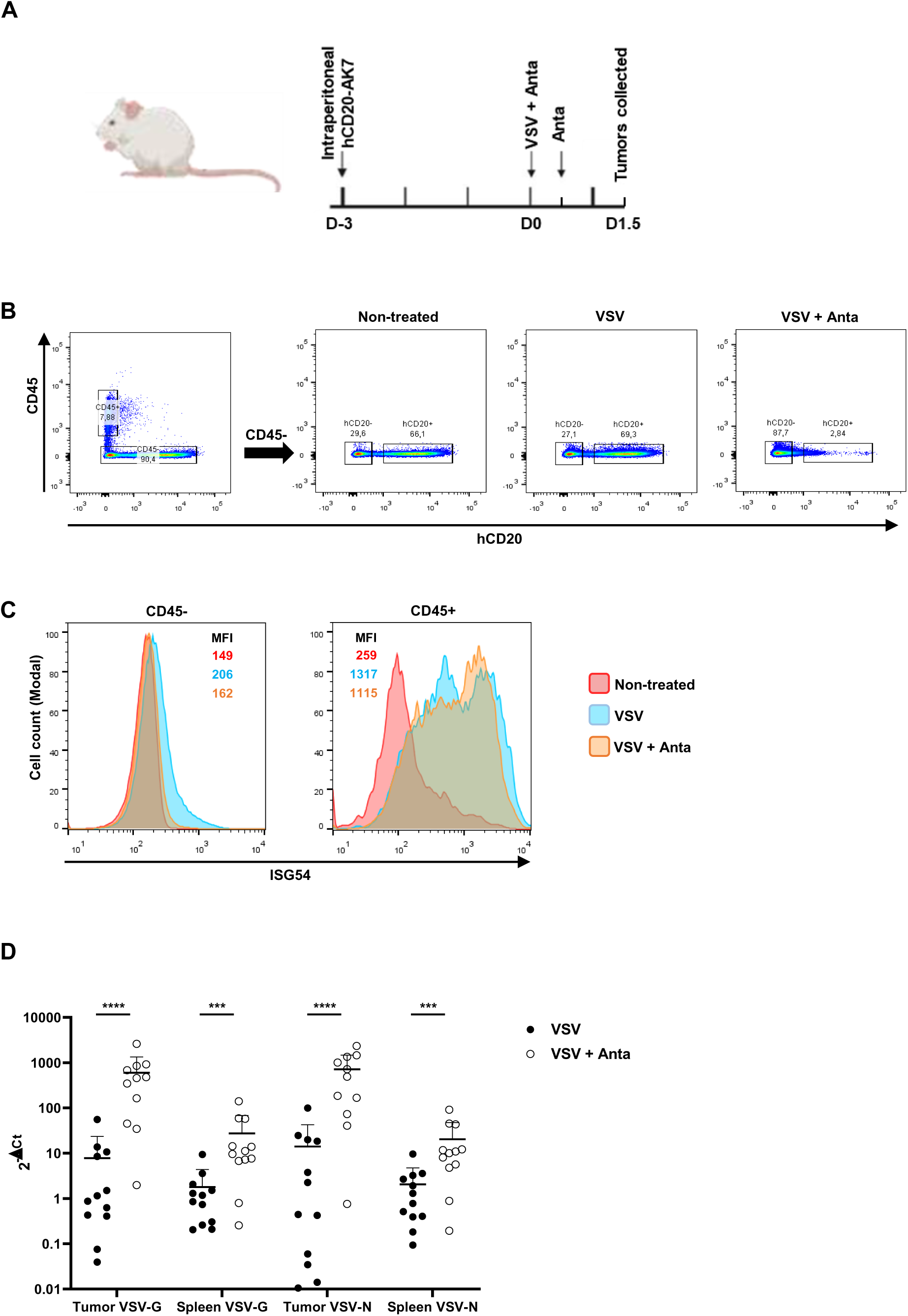
the hCD20-targeted IFNAR1 antagonist reduces the ISG expression and increases VSV replication in hCD20+ tumor cells *in vivo*. (A) hCD20+ AK7 tumor bearing mice were injected IV with 1.5×10^7^ PFU of VSV alone or simultaneously with 10 μg of the hCD20-targeted IFNAR1 antagonist (Anta) by intraperitoneal injection followed by a second dose of 10 μg of the antagonist 12 h later. Tumors and spleens were retrieved 24h later. B) The expression of hCD20 on CD45- tumor cells was measured by flow cytometry. C) The expression of *ISG54* was measured in CD45-and CD45+ cells from dissociated tumors by flow cytometry. D) *VSV-G* and *VSV-N* expression were measured by qPCR. ΔCt represents the difference of cycle numbers between the target gene and the endogenous control gene *Rplp0*. Results are the mean ± SEM of three independent experiments. Viral genes expressions are expressed as 2^-ΔCt^. Expression fold change between VSV+anta and VSV are written in red.

### The hCD20-targeted IFNAR1 antagonist reduces ISG expression replication in hCD2 + AK7 tumor in vivo

To ensure that the antagonist inhibits ISG expression induced by VSV in tumors, we performed a complementary analysis to determine the transcriptomic profiles in some of the tumor samples. The transcriptomic analysis was performed using the NanoString technology with two different panels: «Myeloid Innate Immunity» (754 genes) and «Pancancer Immune Profiling» (770 genes). We found that VSV and the VSV/antagonist combination activates mainly the antiviral pathways, especially the IFN-I response, in tumors and in the spleen (Supplemental figure 3) compared to untreated mice. When comparing mice treated with the VSV/antagonist combination to those treated with VSV alone, we observed a markedly reduced induction of the IFN-I response in the tumor (Figure 6). Importantly, we observed within the tumor a strong decrease in the expression of ISG classified according to the Gene Ontology-Biological Processes (GO-BP) under the “response to IFN-I” pathway (Figure 6C). We also noticed a decrease in the expression of other genes that are not part of the leading-edge genes in the “response to IFN-I” GO-BP pathway but are nevertheless well-known ISG such as *CD274, Cmpk2, Psmb10, Ifit1, Rsad2, Tapbp, Ddx58, Stat2, Psmb8, Ifi35, Herc6, Xaf1, Ifit3, Nlrc5, Tap1, Isg20 and Mx1* (Figure 6A) ^19^. We also observed a distinct transcriptomic profile in the spleen in the presence of the antagonist (Supplemental figure 4A-4B) that differs from that observed in the tumor, both quantitatively and qualitatively (Figure 6A–6C). We notably observed modulation of several growth factor signaling pathways, including macrophage colony-stimulating factor and platelet-derived growth factor signaling, which were upregulated in the spleen in the absence of the antagonist (Supplementary Figure 4C). These differences might account for different inflammatory profiles and viral loads in both tissues as well as to the reduced IFNAR signaling in the tumor upon antagonist administration. Importantly, only a very week reduction of the IFNAR signaling was observed in the spleens of mice treated with the VSV/antagonist combination versus mice treated with VSV alone (Supplemental figure 4C) as compared to tumor samples (Figure 6C), demonstrating that the antagonist acts preferentially in the tumor.

**Figure 6:**
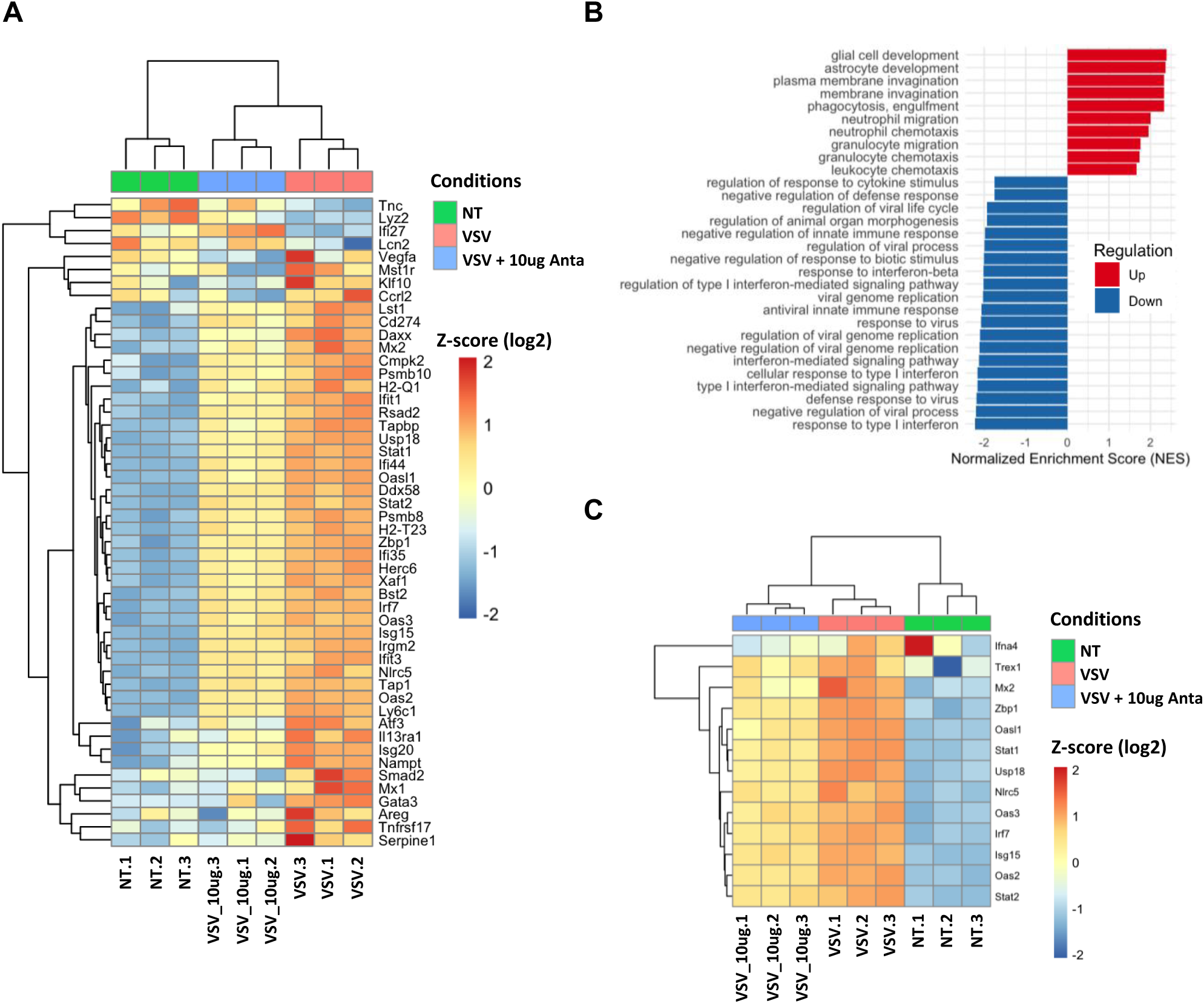
The expression of ISG is induced in the tumor from mouse treated with VSV and is decreased when VSV is combined with the tumor IFNAR1 antagonist. Tumors were implanted by intraperitoneal injection of 5×10^6^ hCD20+ AK7 cells and 3 days later, mice were injected with either intravenous PBS, intravenous 1.5x 10^7^ PFU of VSV or intraperitoneal 10μg of targeted IFNAR1 antagonist (Anta) along with intravenous 1.5×10^7^ PFU of VSV, followed by a second dose of 10 μg of the antagonist 12 h later. After 36 hours after VSV injection, tumors and spleens were harvested. Transcriptome analysis of tumors was performed by nanostring using the «Myeloid Innate Immunity» (754 genes) and the « Pancancer Immune Profiling» (770 genes) panels. Genes expression from both panels were merged and results from genes present in both panels (346 genes) were averaged. A) Heatmap of the top 50 genes between VSV/antagonist vs VSV ranked by limma t-statistic. Colors scale represent z-scores of log2-normalized expressions across samples. B) Pathway analysis using GO-BP. C) Heatmap of the leading-edge genes from the “response to type I interferon” pathway.

### The combination of VSV and the hCD20-targeted IFNAR1 antagonist slightly increases survival of mice bearing hCD20+ AK7 tumor

We next tested the therapeutic effect of the VSV/antagonist combination in mice bearing hCD20+ AK7 tumors (Figure 7A). In an initial experiment aimed at evaluating mouse survival, we observed signs of neurotoxicity (e.g. hind legs paralysis) in hCD20+ AK7 tumor-bearing mice several days after receiving the hCD20-targeted IFNAR1 antagonist and 1.5×10^7^ PFU of VSV (Figure 7B). We hypothesized that the hCD20+ AK7 tumors treated with both VSV and the antagonist could act as a replication reservoir for the virus and amplify the global viral charge to levels associated with neurotoxicity. Hence, in a second experiment, we reduced the dose of VSV to 1.5×10^6^ PFU. In this setting, we observed a trend (p = 0.0513) toward increased survival in mice receiving the VSV/antagonist combination treatment compared with those treated with VSV alone (Figure 7C).

**Figure 7:**
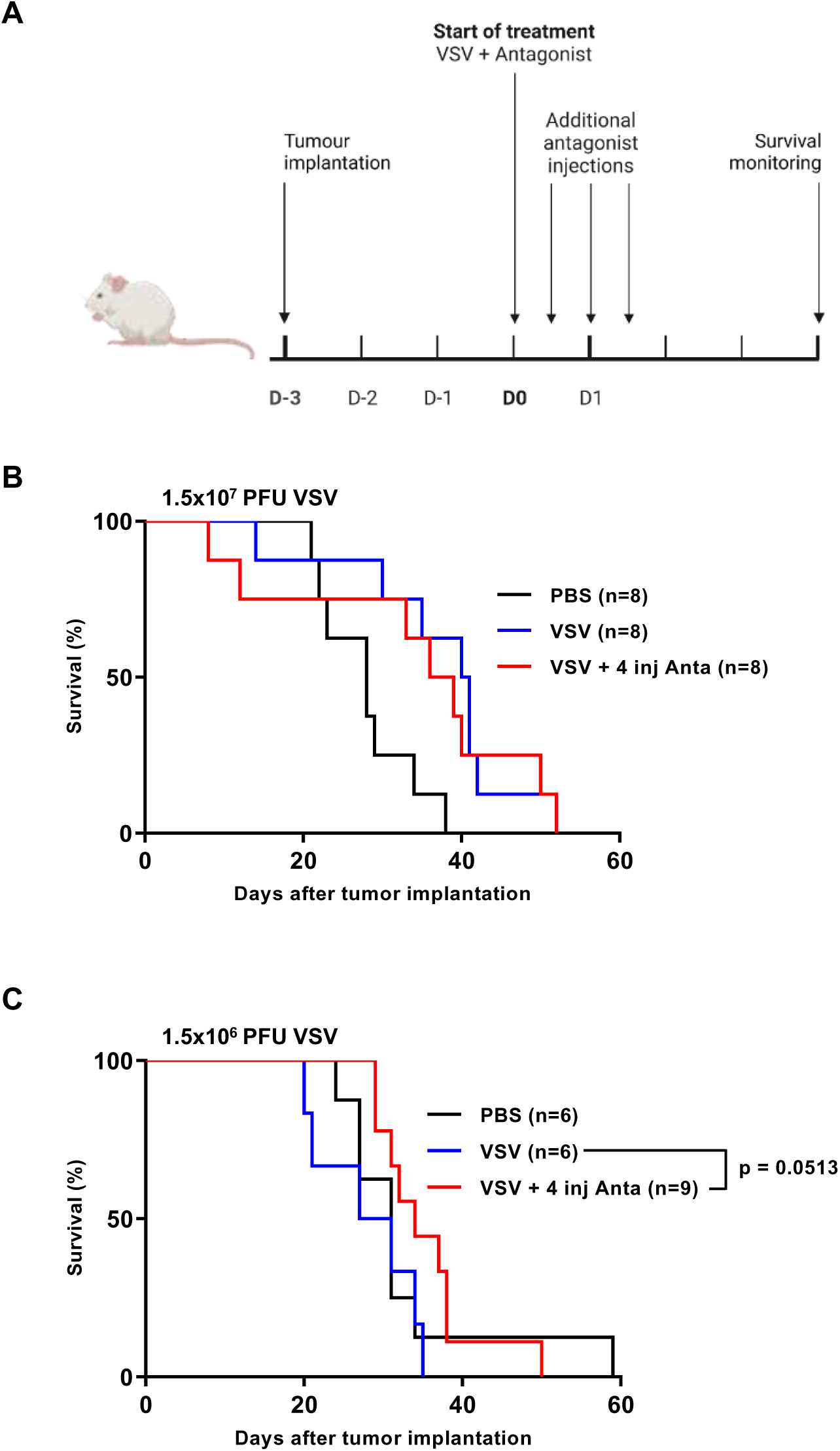
the combination of VSV and the hCD20-targeted IFNAR1 antagonist slightly increases survival of hCD20+ AK7 tumor bearing mice. A) Timeline of the experiment. Tumors were implanted by intraperitoneal injection of 5×10^6^ hCD20+ AK7 cells. 3 days later, 1.5×10^7^ PFU (B) or 1.5×10^6^ PFU (C) of VSV were injected intravenously alone or in combination with an intraperitoneal injection of 10 μg of the hCD20/mIFNAR1 antagonist. 3 other injections of the antagonist 12h apart were performed (4 injections in total). Mice survival was then monitored.

## Discussion

In this study, we report a novel cell-type specific IFNAR inhibitor with strong efficacy. We show *in vitro* and *in vivo* that it is possible to block IFNAR signaling in a particular cell type using an antagonist formed of a conjugate of two nanobodies specific for both a cell-specific marker and IFNAR1. We demonstrate in an *in vivo* setting that this antagonist can be used to block IFNAR signaling specifically in B lymphocytes, DC and mouse-engrafted tumor cells. In addition, we show that in mice treated with oncolytic VSV, the antagonist allows a better replication of the virus in tumors.

After tumor dissociation, flow cytometry analysis showed that IFNAR signaling is not blocked by the hCD20-targeted IFNAR1 antagonist in CD45+ non-malignant cells from the tumor, contrary to CD45- tumor cells. This result suggests that IFNAR signaling is preserved on non-malignant cells, especially immune cells. This is of particular importance since IFN-I plays a major role in the induction of the anti-tumor immune response, in particular by activating antigen-presenting cells ^13,20,21^ or NK cells ^22^. Thus, the tumor cell-targeted IFNAR1 antagonist should preserve IFNAR signaling in these immune cells allowing to keep their involvement in the induction of an antitumor response.

Surprisingly, the improved VSV replication in tumors of animals treated with VSV combined with the antagonist did not lead to a significant increase of the survival suggesting that other factors besides IFNAR signaling are able to limit VSV oncolytic activity *in vivo*. Furthermore, the replication and the survival experiments are difficult to compare, as the increased VSV replication was observed 24h after we treated the mice with 1.5×10^7^ PFU and the antagonist. However, this dose of VSV was associated with toxicities. We hypothesize that VSV produced by infected tumor cells spill over in the mouse circulation and reach toxicity level. When reducing the VSV dose by 10-fold (1.5×10^6^ PFU), the combination modestly increased mice survival, without any observed toxicity. Since we observed toxicity at high doses of VSV combined with the antagonist, one possibility to reduce toxicity would be to use a VSV or another virus, that is more attenuated such as VSVΔ51 ^27^. This virus is more sensitive to the IFN-I response and should exhibit less neurotoxicity when its replication is increased within the tumor.

Recently, it has been shown that VSV infection of non-cancer cells alone can contribute to tumor regression, independently of replication in tumor cells ^23^. In this study, the level of replication of VSV in tumor cells did not affect its oncolytic efficacy, which is consistent with our findings. In addition, we observe a markedly decrease in the global IFN-I response within the tumor in the presence of antagonist which may affect the antitumoral immune response. We also observe a slightly lower IFN-I response in the spleen of mouse treated with the VSV/antagonist combination compared to VSV alone. This reduced IFN-I signaling in the tumor and, to a lesser extent, in the spleen may affect the antitumor immune response, despite the enhanced viral replication in the tumor induced by the antagonist.

Among possible factors besides IFNAR signaling that might limit VSV oncolytic activity *in vivo*, it is worth mentioning that VSV can induce the secretion of other cytokines/proinflammatory factors that can also affect viral replication, tumor properties and immune responses. Notably, El-Sayes *et al*. showed that VSV triggers the production of IFN-I that induces the expression of PD-L1, which could in turn affect the antitumor immune response ^24^. In agreement with this observation, we observed an upregulation of the PD-L1 encoding gene (*CD274*) in both VSV and VSV/antagonist combination treated mice (Figure 7B). This further suggests that the use of PD-L1 inhibitors could improve OV-based therapeutic approaches ^25^. El-Sayes *et al.* also showed that the combination of anti-IFNAR antibodies and VSV is as efficient as the combination of anti PD-L1 and VSV to increase mouse survival ^24^. Further studies are required to better understand this discrepancy between viral replication and the weak survival benefit.

Worthy of note, OVs can be engineered to produce antibodies and antibody-derived molecules to enhance antitumor responses ^4^. In addition, OVs can also be armed with immunostimulating cytokines to locally boost the immune response. In this context, the enhanced viral replication mediated by the antagonist should increase the production of the therapeutic protein in the tumor thereby improving the efficacy of armed VSV. Our work could also have implication for the improvement of CAR-T cell therapies in which OV-derived IFN-I restricts CAR-T cell therapy ^26^. Thus, an antagonist that targets the IFNAR on CAR T cells may be able to restore their efficacy.

Finally, the duration of IFNAR neutralization may be a key point in this type of strategy. The minimum time of VSV replication in the tumor to induce efficient rejection should be further studied, in particular tumor cell lysis and the release of sufficient amounts of damage-associated molecular patterns (DAMPs) molecules to enable an effective immune response^28^. However, one must also be cautious about the priming of a strong antiviral immune response that may hinder the efficacy of viral oncolytic therapy^29^. Further work is needed in order to verify if the limited effectiveness of our combinatory therapy is due to a potent antiviral immune response.

Overall, our work demonstrates the *in vivo* efficacy of a novel cell type-specific IFNAR antagonist. Such antagonists are interesting tools to study *in vivo* the contribution of IFNAR signaling on different cell types, especially immune cells, in the context of infection or cancer. They would be time-saving and cost-effective alternatives to mouse engineering for cell type-specific IFNAR knock downs. These antagonists can also be used in parallel of cell-targeted activators of IFNAR signaling such as conjugates of a mutated IFN-α2 and nanobodies specific for different cell markers that we previously produced and that allow to turn on IFNAR signaling in specific cell types ^12–14^. This array of innovative tools able to specifically block or to specifically induce IFNAR signaling have multiple applications, both for addressing fundamental questions in immunology and for advancing innovative therapeutic approaches.

## Materials and methods

### Tumor cell lines

hCD20+, mCD20+ and unmodified B16 mouse melanoma cell lines were cultured in DMEM culture medium supplemented with 10% FBS, glutamine and penicillin/streptomycin ^14^. hCD20+ and hCD20- AK7 cell lines were cultured in RPMI medium supplemented with 10% FBS, 2 mM glutamine and 100 U/ml penicillin/ 100 µg/ml streptomycin. hCD20+ AK7 clones were obtained by transfection of an hCD20 expressing plasmid ^14^ and selection with 300 µg/ml of G418. A sub-clone stably expressing hCD20 when implanted *in vivo* was obtained after 18 days of growth in C57/BL6 mice.

### VSV production and purification

VSV (Indiana strain) was produced and purified as previously described ^30^. Briefly, BHK cells were seeded and transfected with a plasmid encoding the VSV genome and three accessory plasmids coding for the N, P and L viral proteins. Cells were then infected with a Vaccinia Virus coding for T7 for 1 hour. The supernatant of the cells was collected 24h later and filtered at 0.2 µm. VSV was then amplified on BHK cells for 24h. Then, the supernatant containing the VSV particles is ultracentrifuged at 100,000 g for 1h on a 10% sucrose PBS cushion twice. The viral particles were titrated by plaque forming assay and stored at -80°C.

### Targeted IFNAR1 antagonist production and purification

Lamas were immunized against the ectodomain of a His-tagged IFNAR1 subunit. Peripheral blood was collected and the total RNA of lymphocytes was extracted and used as a template for cDNA synthesis. The obtained VHH cDNA library was then screened for IFNAR1-targeting VHHs by phage display screening technique. By recombinant engineering, the anti-IFNAR1 VHHs were fused with a high affinity anti-mCD20 or anti-hCD20 VHH ^13,14^. Both VHHs were connected with either a 20-fold GGS motif linker or a PASylated linker composed of 300 amino acids (15 repeats of the motif PAAPAPSAPAASPAAPAPAS) ^15^.

### Type I interferons

Two types of IFN-I were used: IFNα-11 and a natural mix of IFN-I (IFN-α and IFN-β). IFN-α11 was obtained by protein production from HEK cells transfected with a plasmid coding for a recombinant IFN-α11 with a His-tag and a C86S mutation ^12^. The natural mix of IFN-I was obtained by infecting C243 cells with Newcastle disease virus (NDV) and subsequently purifying the supernatant for IFN-I ^31^.

### Cytopathic effect measurement

10,000 AK7 or 5,000 B16 cells were seeded in 96-well flat bottom plates. The day after seeding, cells are incubated 30 minutes with the targeted antagonist, followed by exposition to 100 U/ml of IFN-α11 in a 37°C incubator. After 24hrs of incubation with the antagonist and/or IFN-α11, tumor cells are exposed to VSV at an MOI of 0.12 for AK7 and 0.6 for B16 cell lines. After 48h of infection, cells were fixed and stained with a solution of 0.125% crystal violet, 3.7% PFA and 0.15 nM NaCl for 10 minutes at room temperature. After drying, crystal violet was dissolved in a 0.2% Triton X-100 solution and quantified with a colorimeter at 570nm.

### Animals

Female C57/BL6N and Balb/c mice (Charles River) of 8 weeks of age or more were used in the *in vivo* experiments. All animal experiments were approved by the local ethics committee (project granted by the French Ministery of Research, #27051).

### IFN-I and mCD20-targeted antagonist treatment

Female C57/BL6 or Balb/c mice were injected intravenously by the caudal vein with PBS, 30,000 U of the natural mix of IFN-I or a combination of 10μg of mCD20-targeted IFNAR1 antagonist followed 5 minutes later by 30,000 U of a natural mix of IFN-I. Mice were euthanized 17 h later and the spleens were recovered.

### VSV and hCD20-targeted antagonist treatments

Human CD20+ AK7 cells were collected and washed with cold PBS. 5×10^6^ cells in 100 μl of PBS were then injected intraperitoneally to female C57/BL6 mice. Three days after tumor implantation, mice were treated with PBS, VSV or a combination of VSV and the hCD20-targeted antagonist. 1.5×10^6^ or 1.5×10^7^ PFU of VSV in a 100 µl bolus were injected intravenously through the caudal vein. 10 µg of the targeted antagonist in a 50 µl bolus was administered intraperitoneally with additional injections 12h apart. Following euthanasia of the animals, the spleens and tumors of mice were collected. Blood was collected from isoflurane anaesthetized mice through the retro-orbital sinus capillary and deposited into 1,5 ml tubes at room temperature. After centrifugation, supernatant serums were collected and stored at - 80°C for subsequent IFN-α measuring.

### IFN-α detection by Enzyme-Linked Immunoabsorbent Assay

The presence of IFN-α in mice sera was quantified by ELISA (PBL) according to manufacturer instructions. Samples were loaded into pre-coated 96-well plates and treated with proper reagents. Absorbance was then measured with a plate reader at 450 nm. Sample results were interpolated with a standard curve of IFN-α provided by supplier.

### Tumor and spleen dissociation

AK7 mesothelioma tumors were dissociated by an enzymatic (Gibco Collagenase I and IV) and mechanical procedure. Dissociated tumor and splenic cells were then incubated with a Fixable Viability Stain 450 (Invitrogen) viability marker for 30 minutes, on ice, protected from light. The splenocytes were next incubated with anti-mCD16/CD32 Fc Block antibodies (BD), to avoid Fc receptor-dependent binding from the flow cytometry labelling antibodies.

### Flow cytometry analysis

For extracellular staining, cells were labelled with various antibodies: PE-labelled anti murine PD-L1 (Biolegend), APC-H7 labelled anti human CD20 (BD Biosciences) or PE-labelled anti murine CD45 (BD Biosciences). After extracellular staining, cells were fixated for 30 minutes at room temperature with IC Fixation Buffer (InVitrogen). After fixation, cells were subjected to two cycles of washes with Permeabilization Buffer (Invitrogen). Samples were then labelled with rabbit polyclonal anti-ISG54 primary antibody (Invitrogen) and subsequently with an Alexa 488-labeled goat anti rabbit IgG secondary antibody (Invitrogen). In the case of pSTAT1 detection, cells were fixated with Phosflow Fix Buffer 1 (BD Biosciences) and permeabilized with Phosflow Perm Buffer III (BD Biosciences). Cells were then stained with a PE-labelled anti-pSTAT1 (Y701) antibody (BD Biosciences). Fluorescence signal was analysed with a BD FACS Canto II flow cytometer. Data were analyzed using FlowJo Software.

### Immunoblotting

A BCA assay was used to quantify the concentrations of protein in the samples (Interchim). The PVDF membranes containing the samples were incubated with a solution of 0.1% TBS Tween (TBS-T) and 5% milk powder for 1 hour at room temperature, under slow agitation. The membranes were then incubated with the primary antibody (anti-ISG54 Invitrogen; anti-βactin R&D) diluted in 0.1% Tween 5% milk TBS solution. Membranes were incubated with an HRP enzyme-coupled secondary antibody solution (both anti-mouse and anti-rabbit from Interchim) diluted in 5% milk TBS. Membranes were incubated with Clarity Western ECL Substrate Revealing Kit (Bio-Rad). After incubation luminescence was read with a Bio-Rad Chemidoc MP Bio-Imager. Two channels were used in the imaging: a chemiluminescence channel to detect the light emitted by the enzymatic activity of the HRP, and a second colorimetric channel necessary for the visualization of the size ladder.

### Quantitative polymerase chain reaction analysis

A Macherey-Nagel Nucleospin RNAplus Extraction kit was used for RNA extraction of the samples according to manufacturer instructions. The integrity and concentration of the RNA samples were assessed with Agilent Nano 6000 chips (Agilent). RNA samples were used to synthesize cDNA for the qPCR analysis using an M-MLV reverse transcriptase (New England). A qPCR was performed on previously obtained cDNA samples using a Maxima Sybr Green Master Mix (ThermoFisher). Samples were analyzed for different gene expressions using primers designed specifically for this project (*VSV-G*: fw-TGCCCGTCAAGCTCAGATTT, rv-AGCATGACACATCCAACCGT; *VSV-N*: fw-TGTGCCTCGTTCAGATACGG, rv-CGGTTCAAGATCCAGGTCGT). The qPCR analysis was performed using a QuantStudio 3 Real-Time PCR System (ThermoFIsher) at adequate T_m_ settings for our primers. The quantification of gene expression was normalized with Rplp_0_ expression as the endogenous control.

### Transcriptomic study

RNA expression profiling of tumor and spleen cells was performed using nCounter® Mouse Myeloid Innate Immunity V2 Panel (754 genes) and nCounter® Mouse PanCancer Immune Profiling panel (770 genes). Gene expression analysis was conducted with the NanoString technology. Briefly, 5 μL/sample containing of 50 ng of total RNA was combined with the nCounter® reporter CodeSet (3 μL) and nCounter® capture ProbeSet (2 μL) along with hybridization buffer (5 μL) for an overnight hybridization reaction at 65 °C. The reaction was then cooled to 4 °C, and the samples were purified, immobilized on a cartridge, and the data were assessed using the nCounter SPRINT. Normalized genes expression from both panels were obtained from the nCounter analysis software, and merged. Normalized expression of genes present in both panels (n=346) were averaged. After metadata annotation, merged dataset was log transformed and principal component analysis (PCA) performed on the transformed data using the top 50 most variable genes with the prcomp function (stats package, version 4.5.2). Differential expression (limma package, version 3.64.3) and gene set enrichment analyses (GSEA) were performed using the clusterProfiler (version 4.16.0) with minGSSize = 10, maxGSSize = 500, pvalueCutoff = 0.05, using Gene Ontology-Biological Processes (GO-BP) database. Data analysis was done using R studio (version 2026.01.0+392).

### Statistical analysis

For the ISG expression in splenocytes experiments, a non-parametric, two-tailed Mann-Whitney test with an α of 0.05 was performed. For (RT)qPCR experiments that measure the expression of several genes, a non-parametric, one-tailed Mann-Whitney test was performed. For the survival experiments, a non-parametric Mantel-Cox test was performed.

## Data availability statement

All data are available upon reasonable request to the corresponding author.

## Acknowledgements

We thank Gilles Uzé for his invaluable help with this project. We thank the imaging facility MRI (Montpellier), which is part of the UAR BioCampus Montpellier and a member of the national infrastructure France-BioImaging, supported by the French National Research Agency (ANR-10-INBS-04, “Investments for the future”) as well as Cytocell (Nantes) platform for their expertise and support in flow cytometry analysis. We also thank the animal facility of the Institute for Neurosciences Montpellier, (RAM-Neuro; Montpellier), which is part of the UAR BioCampus Montpellier, and UTE (Nantes) animal facilities. We also thank Cecile Monzo and Carole Crozet that are in charge of the Nanostring Sprint Profiler platform in the INM, for providing assistance in the transcriptomic studies. This research was funded by the “Institut National du Cancer (INCA PLBIO 2021-070)”, “Foundation ARC”, the “Ligue régionale contre le cancer Grand Ouest” (Comité départementaux CD16, CD22, CD44, CD49, CD72, CD79 and CD85), “L’association ARSMESO44”, the “LabEX IGO” (ANR-11-LABX-0016-01) and “MabImprove” (ANR-10-LABX-53-01) programs supported by the National Research Agency via the investment of the future program and Orionis Bioscience. TTO was supported by the “Ligue nationale contre le cancer”.

## Authors contributions

Conceptualization: TTO, MP and JFF; Performed experiments: TTO, YB, MS, JN, GG, CC, VD, SD and MNG; Data analysis: TTO, JN, LT, NB, MP and JFF; Writing-original draft: TTO, MP and JFF; Writing-review&editing: TTO, NB, JT, MP and JFF; Funding acquisition: JFF.

