## supplemental figures for "A novel tumor-targeted interferon-alpha/-beta receptor 1 antagonist increases replication of oncolytic vesicular stomatitis virus in a mouse mesothelioma model"

#### Supplemental figure 1

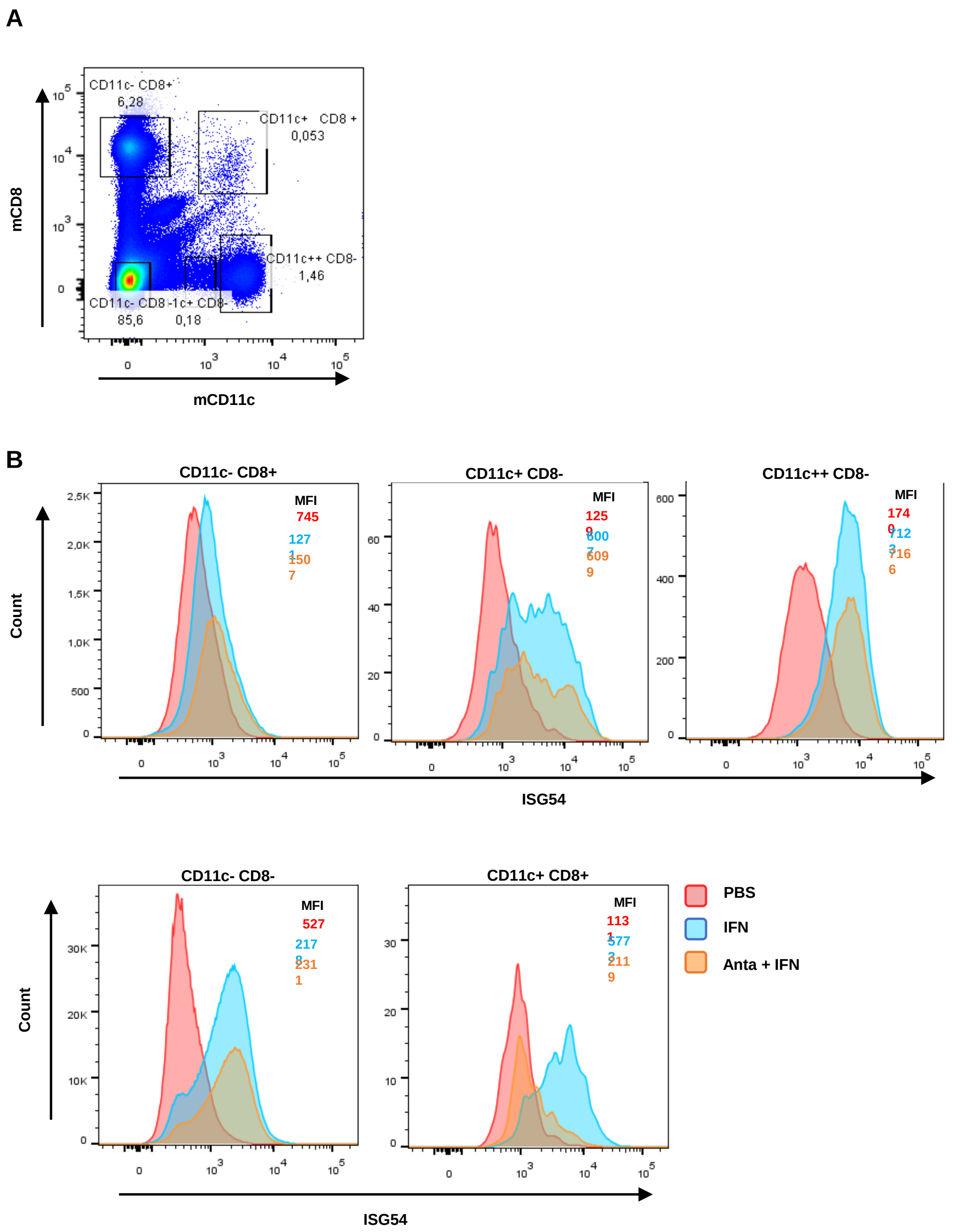


**Supplemental figure 1**: The use of an IFNAR1 antagonist targeted towards CLEC9A-expressing cells can neutralize IFN-induced ISG54 expression *in vivo* in CLEC-9A+ CD11c+ CD8+ dendritic cells. DBA/2 mice were injected intravenously with either PBS, 30000 U of a natural mix of IFN I or 30 µg of the CLEC9A-targeted antagonist followed by a natural mix of IFN I, 5 minutes later. Spleens were recovered 24h later and the expression of ISG54 was analysed by flow cytometry on several splenocytes populations, among them, the CLEC9A+ splenocytes co-expressing CD11c and CD8. A) gating strategy used to identify cell populations. B) histograms represent ISG54 expression on the different cell populations.

#### Supplemental figure 2

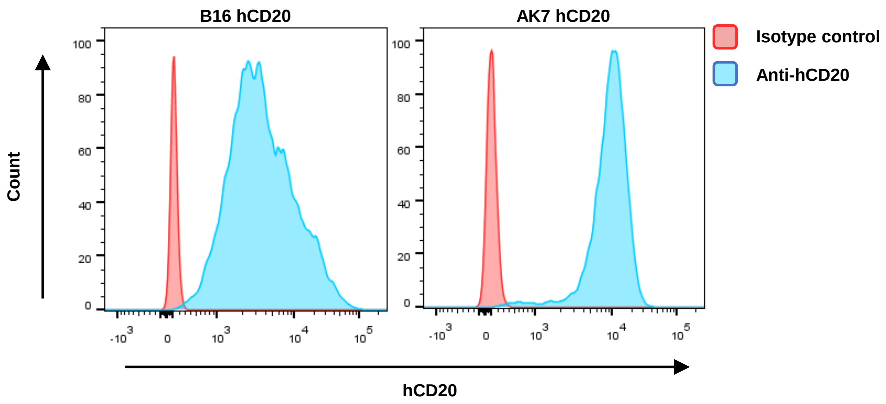


**Supplemental figure 2**: Expression of hCD20 on the hCD20 B16 and hCD20 AK7 cell lines. In vitro cultured cells were stained with an anti-hCD20 APC-H7 labelled antibody or an isotype control antibody and the fluorescence emission was then analysed by flow cytometry.

### Supplemental Figure 3

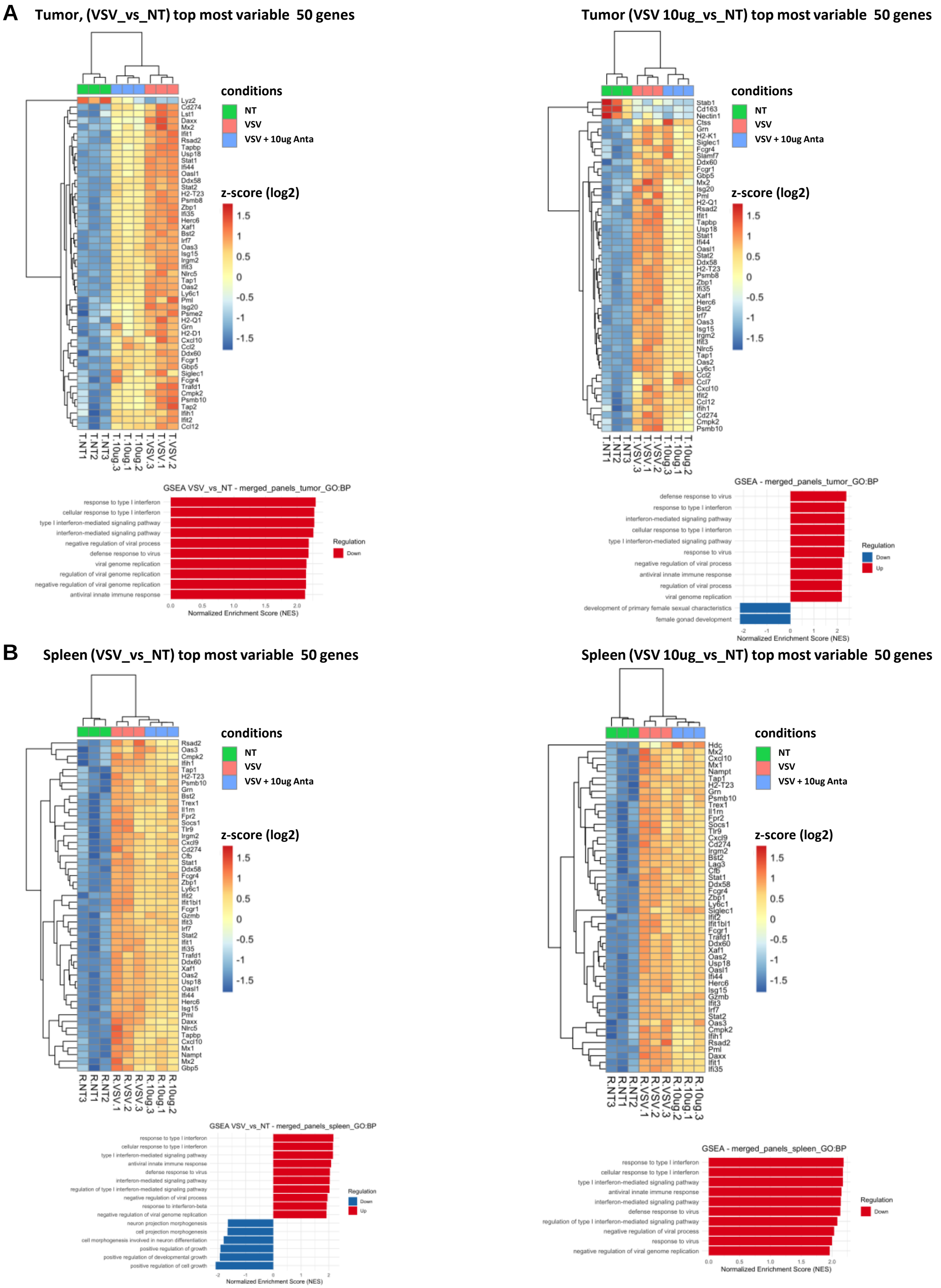


**Supplemental figure 3:** VSV-induced inflammation of the tumor and the spleen. Tumors were implanted by intraperitoneal injection of $5x10^6$ AK7 hCD20 cells and 3 days later, mice were injected with either intravenous PBS, intravenous $1.5x10^7$ PFUs of VSV or intraperitoneal 10µg of targeted IFNAR1 antagonist along with intravenous $1,5x10^7$ PFUs of VSV. 36 hours after VSV injection, tumors and spleens were harvested. Transcriptome analysis of the tumors and the spleens was performed by nanostring using the «Myeloid Innate Immunity» (754 genes) and the « Pancancer Immune Profiling» (770 genes) panels. Genes expression from both panels were merged and results from genes present in both panels (346 genes) were averaged. Heatmaps of the top 50 genes and pathway analysis using Gene Ontology Biological Process (GO-BP) between VSV vs NT and VSV/antagonist vs NT ranked by limma t-statistic in the tumor and in the spleen. Colors scale represent z-scores of log2-normalized expressions across samples. Pathway analysis using Gene Ontology Biological Process (GO-BP).

#### Supplemental Figure 4

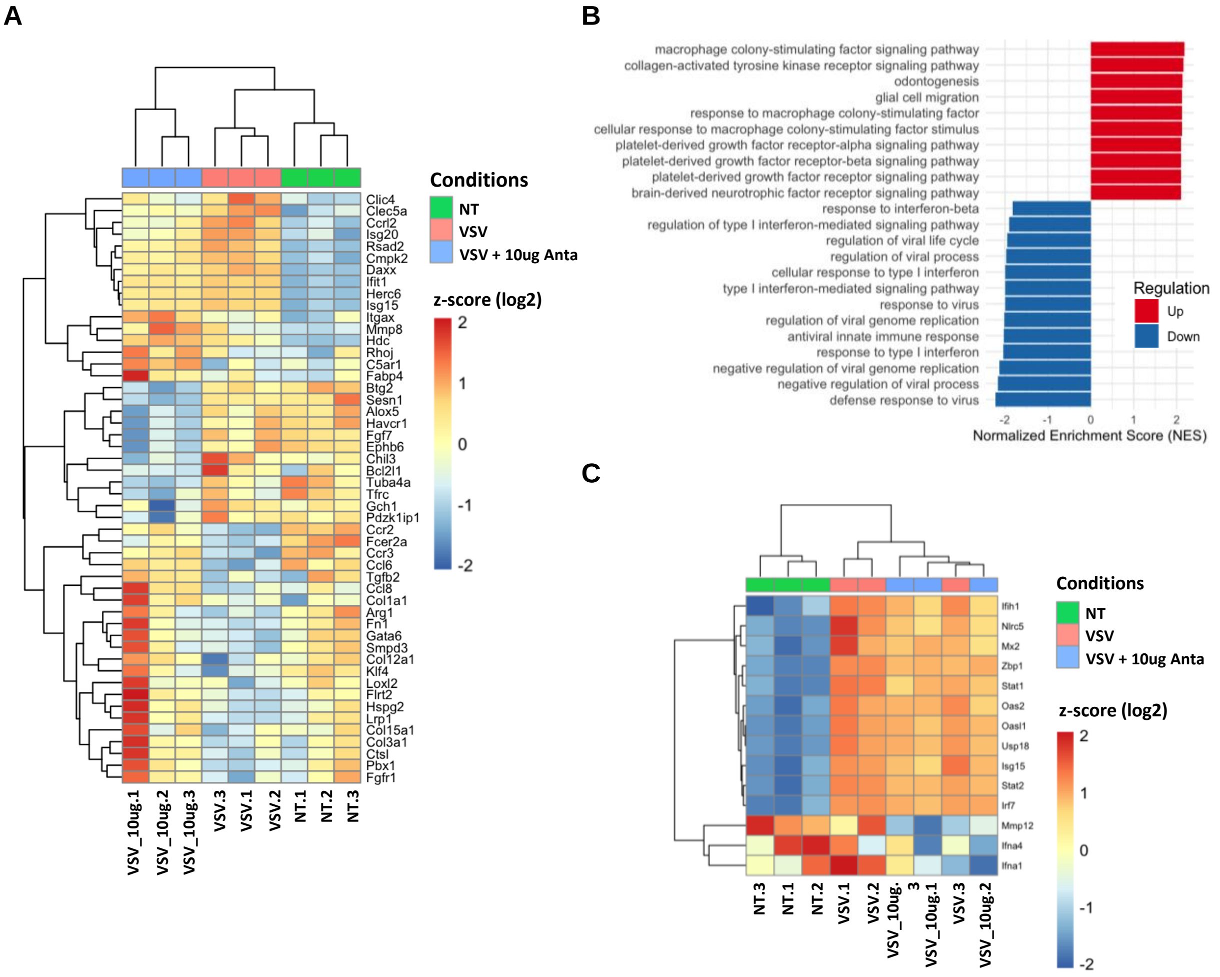


**Supplemental figure 4:** VSV induced IFN I signaling in the spleen that is slightly reduced by the antagonist. Tumors were implanted by intraperitoneal injection of $5x10^6$ AK7 hCD 20 cells and 3 days later, mice were injected with either intravenous PBS, intravenous $1.5x10^7$ PFUs of VSV or intraperitoneal 10µg or 100µg of targeted IFNAR1 antagonist along with intravenous $1,5x10^7$ PFUs of VSV. 36 hours after VSV injection, tumors and spleens were harvested. Transcriptome analysis of spleen was performed by nanostring using the «Myeloid Innate Immunity» (754 genes) and the « Pancancer Immune Profiling» (770 genes) panels. Genes expression from both panels were merged and results from genes present in both panels (346 genes) were averaged. A) Heatmap of the top 50 genes between VSV/antagonist vs VSV ranked by limma t-statistic. Colors scale represent z-scores of log2-normalized expressions across samples. B) Pathway analysis using Gene Ontology Biological Process (GO-BP) database. C) Heatmap of the leading-edge genes from the “response to type I interferon” pathway.
